# Towards estimating marine wildlife abundance using aerial surveys and deep learning with hierarchical classifications subject to error

**DOI:** 10.1101/2023.02.20.529272

**Authors:** Ben C. Augustine, Mark D. Koneff, Bradley A. Pickens, J. Andrew Royle

**Affiliations:** U.S. Geological Survey, Eastern Ecological Science Center, 12100 Beech Forest Road, Laurel, Maryland, 20708 USA; Division of Migratory Bird Management, U.S. Fish and Wildlife Service, 69 Grove Street Extension, Orono, Maine, 04469 USA; Division of Migratory Bird Management, U.S. Fish and Wildlife Service, 11510 American Holly Drive, Laurel, Maryland, 20708 USA

**Keywords:** marine wildlife, abundance estimation, misclassification, CNN,k aerial survey

## Abstract

Aerial count surveys of wildlife populations are a prominent monitoring method for many wildlife species. Traditionally, these surveys utilize human observers to detect, count, and classify observations to species. However, given recent technological advances, many research groups are exploring the combined use of remote sensing and deep learning methods to replace human observers in order to improve data quality and reproducibility, reduce disturbance to wildlife, and increase aircrew safety. Given that deep learning detection and classification are not perfect and that statistical inference from ecological models is generally very sensitive to misclassification, we require study designs and statistical models to accommodate these observation errors.

As part of an ongoing effort by the U.S. Fish and Wildlife Service, Bureau of Ocean Energy Management, and U.S. Geological Survey to survey marine birds and other marine wildlife using digital aerial imagery and deep learning object detection and classification, we developed a general hierarchical model for estimating species-specific abundance that accommodates object-level errors in classification. We consider hierarchical deep learning classification at multiple taxonomic levels subject to misclassification, hierarchically-structured human validation data subject to partial and erroneous misclassification, and an image censoring process leading to preferential sampling. We demonstrate that this model can estimate species-specific abundance and habitat relationships without bias when the assumptions are met, and we discuss the plausibility of these assumptions in practice for this study and others like it.

Finally, we use this model to demonstrate the relevance of the features of the ecological systems under study to the classification task itself. In models that couple the ecological and classification processes into a single, hierarchical model, the true classes are treated as latent variables to be estimated and are informed by both the classification probability parameters and the ecological parameters that determine the expected frequencies of each class at the level the data are being modeled (e.g., site or site by occasion). We show that ignoring the expected frequencies of each class (when they are imbalanced) can cause correction for misclassification to produce biased parameter estimates, but coupling the ecological and classification models allows for the variability in relative class frequency across space and time due to ecological and sampling conditions to be accommodated with spatial or temporal covariates. As a result, bias is removed, classification is more accurate, and uncertainty is propagated between the ecological and classification models. We therefore argue that ability of deep learning classifiers, and classifiers more generally, to produce reliable ecological inference depends, in part, on the ecological system under study.

## Introduction

Aerial surveys are a standard method for estimating the abundances, population trajectories, and habitat associations of a diverse range of wildlife species. Concern over population declines of many species, including species of sea ducks (Bowman et al., 2015), seabirds (Croxall et al., 2012; Paleczny et al., 2015) and marine mammals (Reynolds III et al., 2009), have led to increased interest in population monitoring to aid in assessing the effects of harvest (Koneff et al., 2017) energy extraction (Fox and Petersen, 2019; Ronconi et al., 2015), by-catch (Žydelis et al., 2013), plastics and debris (Wilcox et al., 2015), and other factors. Low-level aerial surveys conducted by human observers have been a cornerstone of waterfowl and marine mammal management in North American since the 1950s (Smith, 1995; Johnson et al., 1997; Zimpfer, 2021), because they have provided the only cost-efficient and feasible means to monitor vast and inaccessible regions. Aerial surveys have also become increasingly important in monitoring near-shore populations of marine wildlife and provide data to assess and mitigate the effects of human activities in marine environments (Palka et al., 2017, 2021).

Traditionally, these surveys are conducted by flying aerial transects or other sample units at low altitude with human observers in the aircraft that locate, count, and classify marine wildlife to species. If conducted without error, these data are appropriate for unbiased estimation of the abundance, distribution, habitat associations, and population trajectories of marine species. However, survey data collection errors that can bias inference are common (Thompson, 2002; Williams et al., 2002; Clement et al., 2017; Brack et al., 2018; Davis et al., 2022), limiting our ability to monitor marine wildlife reliably.

Davis et al. (2022) categorize the possible data collection errors for aerial surveys as 1) nondetection, 2) counting, and 3) misclassification. Nondetection errors occur when an observer fails to detect an individual or group that is otherwise available for detection (i.e., detection probability < 1). Counting errors occur when an observer identifies a group, but incorrectly enumerates the number of individuals in the group. Misclassification occurs when an observer incorrectly records the species of an individual or group. Imperfect detection in aerial surveys has traditionally been accommodated using double sampling (Smith, 1995), double observers (Koneff et al., 2008), or distance sampling (Buckland et al., 2001), particularly mark-recapture distance sampling (MRDS) that uses multiple observers to estimate the probability of detection at a distance of 0 from the transect (Borchers et al., 1998). These models are appropriate when nondetection events are independent; however, in aerial surveys of groups, some forms of nondetection may affect individuals independently, while others may affect the group as a whole (e.g. visibility bias or observer fatigue), making it inappropriate to apply these models to individual-level detections (Davis et al., 2022). Clement et al. (2017) provide a model for imperfect detection at both the individual and group-level by combining MRDS with an N-mixture model (MRDS-Nmix; Royle, 2004), which accommodates the dependence of group-level detections appropriately.

With counting errors, human observers tend to visually underestimate the number of individual animals in a group, and the magnitude of that error typically increases as the group size increases (Erwin, 1982; Frederick et al., 2003; Davis et al., 2022). The MRDS-Nmix model of Clement et al. (2017) accommodates undercounting, but it assumes undercounting results from imperfect detection of individual group members, which are independent (Clement et al., 2017). Independence may not hold in practice if observers are assigning group sizes using mental addition, extrapolation, or approximation (Clement et al., 2017), though other statistical models for miscounting could be used.

Misclassification errors in aerial wildlife population surveys are often not addressed despite the fact that they can induce large biases in wildlife population estimates (Conn et al., 2013; Davis et al., 2022). Attempts to address misclassification bias in population estimation from aerial wildlife survey data typically required costly auxiliary data collection and statistical assumptions that are challenging to meet (Conn et al., 2013, 2014; McClintock et al., 2015). Statistical models to correct for species misidentification have received much attention in the species occupancy literature (Royle and Link, 2006; Chambert et al., 2015; Wright et al., 2020). False-positive occupancy models allow for the estimation of species occupancy probabilities in the presence of false-negatives (non-detection) and false-positives (species misidentification). These models were originally formulated for occupancy data that were “contaminated” with false-positive detections and the inferential goal is to estimate the false-positive rate, used to correct the occupancy parameter estimates for misidentified detections (Royle and Link, 2006). These false-positive occupancy models accommodate “target” and “non-target” classes (species) of detections; however, for multi-species surveys, jointly modeling all species and estimating their pairwise classification probabilities is often more appropriate because false-positives for one species may result in false-negatives for another and joint modeling better accommodates this type of data (Wright et al., 2020; Spiers et al., 2022). Further, species classification can be improved when coupled with an ecological model that jointly estimates the habitat associations and detection characteristics of multiple species simultaneously with the false-positive or pairwise species classification probabilities (“coupled classification”; Kéry and Royle, 2020; Spiers et al., 2022). Regardless of the approach, false-positive occupancy models require either known-identity validation samples (or site occupancy states) or informative priors for the classification probabilities (Royle and Link, 2006; Chambert et al., 2015; Wright et al., 2020).

Aerial surveys of waterfowl or marine mammals do not readily fit within an occupancy modeling framework for which false-positive methods are well-developed because there is typically no replication of survey segments that could be treated as “sites”. Species misclassification could potentially be modeled in a distance sampling framework, but obtaining known-identity validation samples to compare with the human classifications made in flight is difficult and likely cost-prohibitive. More recently, aerial surveys of wildlife populations are being conducted through remote sensing (e.g., optical or thermal sensors) with deep learning used to detect animals in images and classify them to species(e.g., Maire et al., 2015; Kellenberger et al., 2018; Guirado et al., 2019; Hong et al., 2019; Edney and Wood, 2021; Kellenberger et al., 2021). Data collection through remote sensing has the potential to better accommodate misidentification errors because the aerial images provide a permanent record of the surveyed sample unit and validation data can be generated by humans for a subset of all objects classified by deep learning. Therefore, aerial imagery processed by deep learning may provide a more robust sampling framework for marine wildlife as long as non-detection and counting errors as well as any new errors or biases resulting from the deep learning data processing pipeline data are rare or can be accommodated.

Recently, the US Fish and Wildlife Service (USFWS), Bureau of Ocean Energy Management (BOEM), and US Geological Survey (USGS) have begun conducting aerial surveys of this type and training object detection and classification deep learning models with the goal of estimating marine wildlife abundance and distribution patterns (Figure 1). Three unique features of the data processing pipeline are 1) a hierarchical classification scheme with deep learning algorithms applied at multiple taxonomic levels, 2) preferential sampling (Conn et al., 2017; Pacifici et al., 2016) where images unlikely to have objects of interest are excluded from classification at the lowest taxonomic level, and 3) human validation data with hierarchical classifications that may be erroneous at one or more taxonomic levels and may be incomplete (e.g., an object may be identifiable to genus, but not species). Here, we develop a model to accommodate these three features in order to estimate the abundance and habitat associations of sea ducks and other marine wildlife without bias for this system. We then use this model to demonstrate the importance of the ecological model for the classification task itself.

**Figure 1.**
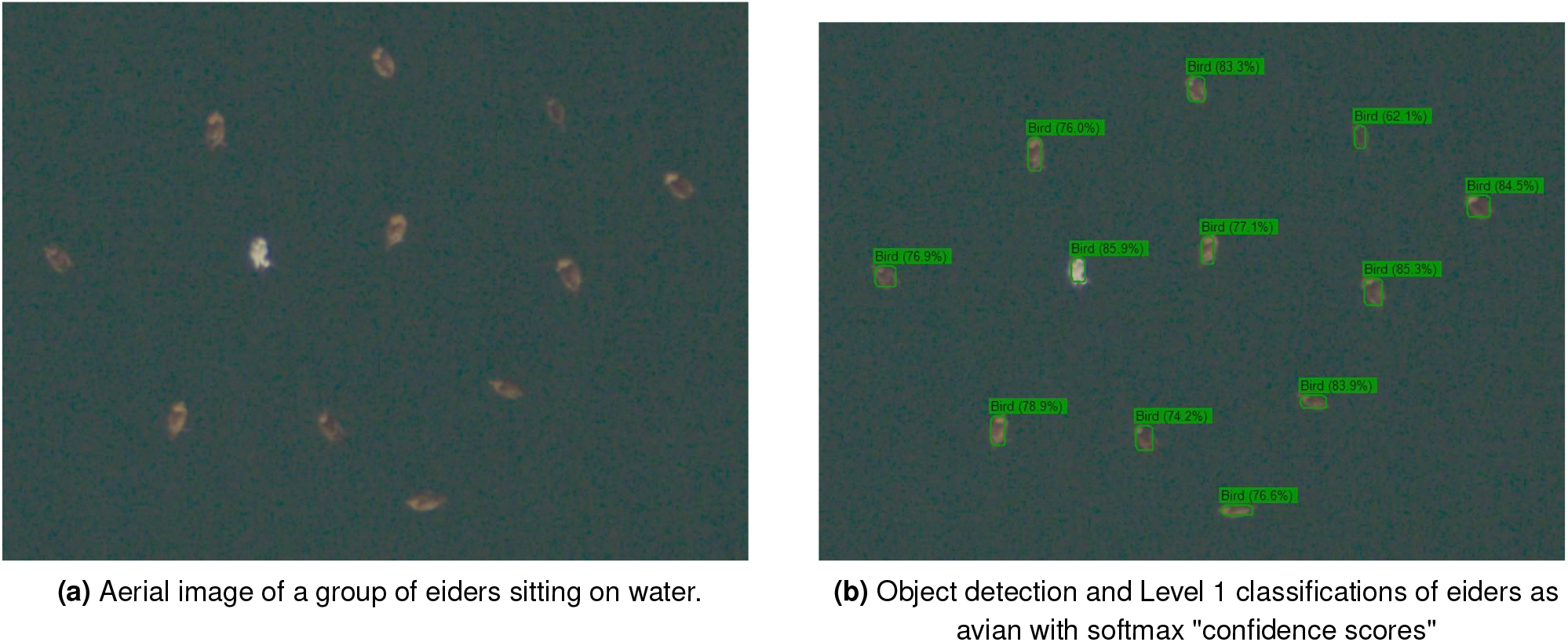
Example aerial imagery, object detection, and Level 1 classification.

## Methods

### Methods – Model Overview

We consider the case where an aerial survey is conducted and 1) the resulting images are subjected to an automated deep learning object detection algorithm (e.g., You Only Look Once (YOLO); Redmon et al., 2016) to identify objects in the images and classify them at a coarse taxonomic scale (e.g., avian, non-avian wildlife, human-made object), 2) the deep learning “confidence scores” (i.e. the softmax vector, Guo et al., 2017) for these coarse taxonomic classifications are used to select images likely to contain objects of interest for classification at a higher taxonomic resolution, and 3) the objects in these selected images are classified to species (or higher taxonomic category) using a convolutional neural net (CNN, LeCun et al., 1998). Because deep learning models are not perfect classifiers, this process introduces classification errors for the objects in the images at both higher and lower taxonomic scales, which can bias ecological inferences (Royle and Link, 2006). Further, the purpose of image censoring in Step 2 is to discard the majority of images that do not contain objects of interest (e.g., unoccupied waters) to reduce the computational resources required for both CNN classification and ecological parameter estimation. However, this image censoring process is a case of preferential sampling, where the sampled locations are conditionally dependent on the process of interest (abundance), requiring us to model the censoring process for unbiased inference (Diggle et al., 2010; Pacifici et al., 2016; Conn et al., 2017).

Our solution to the issues outlined above involves the use of hierarchical classification–the object detection algorithm assigns coarse taxonomic categories and the CNN assigns lower, more specific taxonomic categories. Further, we will require human classifications to use as validation data, but humans are often only able to provide higher-order or coarse classifications for many detected objects in this system. To accommodate these features of the data classification process, we introduce what we call a “classification hierarchy” (Table 1), describing the nested relationships between lower and higher-order taxonomic classifications used in the analysis. We define Level 1 to be the highest taxonomic order with *n*^*L*1^ categories, and this is the level at which the object detection algorithm assigns classifications. The lowest-level classification is the level at which the CNN assigns classifications and the level at which we can make inference about abundance and spatial distribution. Usually, the lowest level will be species; however we may need to use a higher taxonomic order, such as genus, for species that the CNN performs poorly at the species level. In our system, the lowest level is Level 3 with *n*^*L*3^ categories. The intermediate Level 2 is only used for the human classification of the validation data in this system, but any number of intermediate levels may be used. Table 1 outlines a hypothetical 3-level classification hierarchy for Atlantic marine wildlife surveys. We define ***A*** to be the enumerated classification hierarchy (Table 1, numeric label) with *α_i,q_* being the value of Level 3 class *i* at hierarchical level *q*, which we will reference below.

**Table 1.** Classification hierarchy containing a hypothetical hierarchical class structure. Level 3 or species is the desired level of inference, with each Level 3 class having a unique row. The enumerated classes are on the left and their meaning is on the right.

| Numeric Label |  |  | Character Label |  |  |
| --- | --- | --- | --- | --- | --- |
| Level 1 | Level 2 | Level 3 | Level 1 | Level 2 | Level 3 |
| 1 | 1 | 1 | avian | eider | common eider |
| 1 | 1 | 2 | avian | eider | king eider |
| 1 | 2 | 3 | avian | scoter | black scoter |
| 1 | 2 | 4 | avian | scoter | white-winged scoter |
| 1 | 2 | 5 | avian | scoter | surf scoter |
| 2 | 3 | 6 | non-avian wildlife | dolphin | dolphin species 1 |
| 2 | 3 | 7 | non-avian wildlife | dolphin | dolphin species 2 |
| 2 | 4 | 8 | non-avian wildlife | turtle | turtle species 1 |
| 2 | 4 | 9 | non-avian wildlife | turtle | turtle species 2 |
| 3 | 5 | 10 | human-made | human-made | human-made |

### Methods – Data Description

We require three sources of data– 1) the censoring outcomes from the object detection algorithm at Level 1, 2) the CNN classifications of the objects in retained images at Level 3 and 3) paired CNN and human validation data to aid in estimating the CNN classification probability matrix (Wright et al., 2020; Spiers et al., 2022). We will describe each in turn. First, we observe a binary censoring outcome for each Level 1 class and each image. For censored images, this is the only data observed. We assume that the censoring outcome at the image level is a function of the total number of Level 1 objects of a specific class or classes in an image; the manner in which censoring is done is flexible and we describe one approach below. Let *z_a,j_* be the binary censoring outcome for Level 1 class *a* in image *j*, with 1 indicating retain and 0 indicating censor. This is the data required to account for nonrandom censoring as it carries information about the number of Level 1 objects in discarded images, though this information is also informative of the Level 3 classes for objects in images that are not censored. Second, the observed data from the retained images, in addition to their censoring outcomes, are object and image-referenced Level 3 classifications. For each survey object *l* located in image *j* we observe a Level 3 classification, 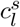 (*s* indicating survey), for all *J* images and *n^s^* objects. The image number of object *l* is indicated by *ρ_l_*.

Third, we have validation data classified by both the CNN and human observers used to inform the CNN Level 3 classification probability matrix estimates. The validation data may come from a subset of the focal survey images, or they may come from a previous survey as long as the CNN is perfectly transferable, by which we mean the CNN Level 3 classification probabilities are exactly the same between the focal survey and the survey that produced the validation data. As with the survey data, each object is extracted and subjected to Level 3 CNN classification. For each validation object *m* = 1,…, *n^v^*, we observe the CNN-predicted Level 3 class, 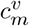. Paired with these CNN classifications, we observe human classifications from multiple observers for these same validation objects. These classifications are hierarchical with three levels as depicted in the classification hierarchy (Table 1). Let 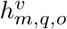 be the human classification for object *m* at hierarchical level *q* from observer *o* for *Q* = 3 levels and *O* = 2 observers. Humans may not always be able to determine the full 3-level classification, in which case, they may classify the first (no classification), 1, or 2 levels. For example, a human may only be able to determine that a object contains an avian or an avian-scoter for a true Level 3 class of black scoter (*Melanitta americana*).

### Methods – Ecological Process Model

In order to accommodate nonrandom censoring and misclassification, we propose a hierarchical model (sensu Kéry and Royle (2020)) with submodels for the ecological process, the Level 1 classification and censoring process, and the Level 3 classification process, which we will describe in turn. In the absence of misclassification, the survey data structured at the object-level consist of the true Level 3 class of each object *l*, 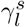, and the associated image number *ρ_l_*. These data can be restructured into Level 3 class by image count data. Let 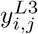 be the count of objects from Level 3 class *i* in image *j*, i.e., 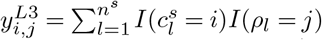, where 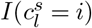 is an indicator function taking value 1 if object *l* contains Level 3 class *i* and 0 otherwise, and *I*(*ρ_l_* = *j*) is an indicator function taking value 1 if object *l* was extracted from image *j* and 0 otherwise. If the CNN is a perfect classifier, then 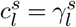 for all *l* and the Level 3 abundance of image-level objects can be directly modeled with a log-linear count model such as Poisson or negative binomial regression which has been done for previous marine wildlife surveys (Zipkin et al., 2014).

We assume the Level 3 class by image counts are well modeled by a negative binomial regression model where the expected number of Level 3 counts in an image varies, perhaps as a function of one or more image-level covariates, which could be features of the transect, date, or habitat. More specifically, 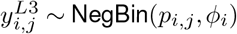. The expected abundance of Level 3 class *i* in image *j* is *λ_i,j_* where log(*λ_i,j_*) = *β*_0,*i*_ + *β*_1,*i*_*x_j_* and *x_j_* is the covariate value for image *j*, *β*_0,*i*_ is the intercept for Level 3 class *i* on the log scale and *β*_1,*i*_ is the covariate coefficient for Level 3 class *i*. Then, we convert the expected count for Level 3 class *i* in image *j* to the negative binomial probability parameter 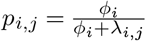 where *ϕ_i_* is the dispersion parameter for Level 3 class *i*.

### Methods – Level 1 Classification and Image Censoring

We accommodate nonrandom image censoring using a model for the censoring process conditioned on the ecological process model. First, the censoring process occurs at Level 1 of the classification hierarchy, so the Level 3 class by image data, ***Y***^*L*3^, must be aggregated to Level 1 to model the censoring process. Let 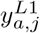 be the Level 1 counts for class *a* in image *j*. Then, 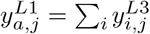 for *α*_*i*,1_ = *a*, e.g., the count of Level 1 avian objects in image *j* is the sum of the counts for the individual avian Level 3 classes. For our application, the in-flight object detection algorithm assigns a vector score of length 3 (avian, non-avian wildlife, human-made) for every detected object and these scores sum to 1. Then, for every image, the maximum score for each of the three Level 1 categories are calculated and the image is retained if any of these scores exceed a threshold score for retention. For example, the decision rule may be to retain images with a maximum avian score of 0.8 or a maximum non-avian wildlife score of 0.9, with no rule for human-made. The general censoring process above leads to the probability of censoring being a function of the total number of Level 1 objects in an image, which we model using logistic regression.

As described above, *z_a,j_* is the observed binary censoring outcome for Level 1 class *a* in image *j*. We assume 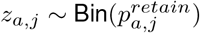, where 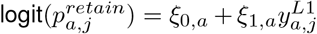. For any Level 1 class *a* that is not of interest, we set 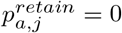, so that *z_a,j_* will always be 0. Then, we retain image *j* if ∑_*a*_*z_a,j_* > 0, i.e., if we pass the retention threshold for any Level 1 class. Alternatively, the relationship between the probability of retaining an image and the number of target objects may be modeled using more flexible functions such as generalized additive models (GAMs; Hastie, 2017). As a second option, instead of modeling binary outcomes (retained, not retained), we could model the censoring process based on the maximum Level 1 scores for each image.

### Methods – CNN and Human Level 3 Classification

The CNN classification process is a model for the CNN classifications we observe, 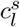, given the true Level 3 class of each survey object, 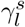. We adopt the classification model used by Wright et al. (2020) and Spiers et al. (2022) for a multi-species occupancy model. Let **Θ**^*c*^ be an *n*^*L*3^ × *n*^*L*3^ classification probability matrix where 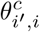 is the probability an object containing Level 3 class *i* is classified as Level 3 class *i*′. Then, the model for the observed CNN classification of object *l* is 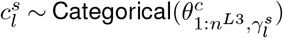. With models for both the Level 3 class by image counts and species misclassification we can jointly estimate the ecological parameters of the abundance regression and the true species of every survey object for retained images (Wright et al., 2020; Kéry and Royle, 2020; Spiers et al., 2022). The final data we require is human validation data in order to aid in the estimation of the CNN species classification probabilities, **Θ**^*c*^.

Typical uses of validation data assume that the validation classifications are complete and without error, that is, humans can supply a Level 3 (species) classification 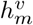 corresponding to the the CNN classification, 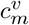 for validation objects *m* = 1 …, *n^v^* without error. Then, using the same misclassification model used for the survey data, we assume 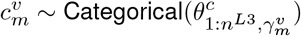. The classification parameters, **Θ**^*c*^, are estimable because ***γ***^*v*^ (the true class) is known with certainty. However, we expect humans interpreting aerial images to make some errors. If there is misclassification error in the human validation data, our estimates of **Θ**^*c*^ will be biased and as a result, so will the estimates of the ecological parameters of interest. A second challenge is that humans will not be able to provide a full classification down to Level 3 for all objects, but may be able to provide a higher-level classification which carries information about the true Level 3 class of an object (e.g., an object classified as scoter, is more likely to be a black scoter than a common eider (*Somateria mollissima*) if human classifiers are generally accurate). This type of partial classification has previously been used by Spiers et al. (2022) for the focal data set subject to misclassification in conjunction with perfect validation data, whereas we will use it for the incomplete and imperfect validation data set itself.

To accommodate imperfect human validation data, we use a double observer model with hierarchical classifications. Double observer designs have been used in traditional aerial surveys to accommodate counting errors made by humans in an airplane (Koneff et al., 2008; Clement et al., 2017) whereas we use a double observer design to accommodate misclassification errors made by humans interpreting objects extracted from aerial images. Because the true classes of the validation data must be estimated, we require a model for the relative frequency of Level 3 classes. For the survey objects, this information at the image level is contained in the parameters for the ecological process model, ***β*** and ***ϕ*** (Kéry and Royle, 2020, Simulation Study 2 and Appendix B below). If validation samples are a subset of the survey data (e.g., Spiers et al., 2022), no additional model for class frequency is required. For simplicity, we assume the validation samples are not a subset of the survey data from which ecological inference will be made, so we do not need to know the ecological and sampling conditions where they were collected. Therefore, we require a model for the relative frequency of Level 3 classes in the validation data set (not structured by image) whose parameters will be estimated in the hierarchical model. For this, we assume 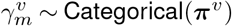 where ***π***^*v*^ is the vector of Level 3 class frequencies for the validation objects.

The final component of the model for human validation data is a submodel for human misclassification. As described above, 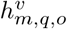 is the human classification for validation object *m* at hierarchical level *q* made by observer *o*. We assume all human classification errors are a function of the true Level 3 class only. The human classification probability matrix **Θ**^*h*^ (h indicates human) is of dimension *n*^*L*3^ × *n*^*L*3^, analogous to the CNN classification probability matrix described above. Similarly, we assume 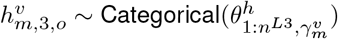, where *h* = 3 corresponds to the Level 3 classification.

To accommodate partial human labels, we assume that hierarchical human classifications are consistent across levels. For example, we assume that if a human assigns an object to be common eider, they necessarily will assign it to be an eider at Level 2 and avian at Level 1. Therefore, if we observe a human Level 2 assignment of eider, the human necessarily would have assigned a Level 3 label of one of the two possible eider species if they were able to make the Level 3 determination. We further assume that the probability a human will be able to assign a full 3-level classification does not depend on the Level 3 class, that is, hierarchical classifications are missing at random with respect to their Level 3 class. For example, the probability a human can only assign an avian-eider classification does not depend on whether the true Level 3 class is common eider or king eider (*S. spectabilis*). A second requirement is that human classifications are missing at random with respect to the CNN classification outcome, e.g., humans are not less likely to provide complete classifications for objects the CNN classifies incorrectly than for the objects the CNN classifies correctly. Finally, we assume the CNN and observed human classifications are independent. We discuss the plausibility of these assumptions in the Discussion.

Given these assumptions that data are missing completely at random (MCAR; Bhaskaran and Smeeth, 2014), we construct the probability model for human classification by marginalizing over the unobserved classifications as follows. First, we calculate the number of human-classified levels for every validation object, 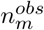, which may take the values 0, 1, 2, or 3. Then, let the probability of the observed vector assignment be

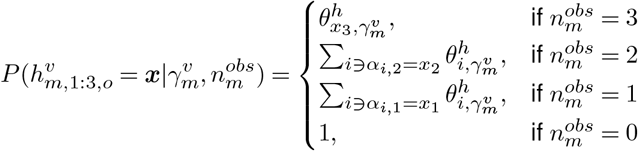

where ***x*** is the observed vector classification assignment of length 3 and *α_i,q_* are indices of the enumerated “classification hierarchy” in Table 1. If *n^obs^* = 1, we sum over the *i* indices of 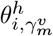 such that *α*_*i*,1_ = *x*_1_. Similarly, if *n^obs^* = 2, we sum over the *i* indices of 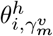 such that *α*_*i*,2_ = *x*_2_.

To give a concrete example, consider the case where the true Level 3 class is black scoter and a human classified it as surf scoter (*M. perspicillata*). Then *n^obs^* = 3, and the probability of this event is 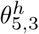; the true species is species 3 and the classified species is species 5. However, if the human can only classify to scoter at Level 2, then *n^obs^* = 2 and the probability of this classification is 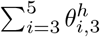, or the summation of the classification probabilities for all the scoters given the true Level 3 class is black scoter. Similarly, if the human can only classify to avian at Level 1, then *n^obs^* = 1 and the probability of this classification is 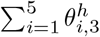, or the summation of the classification probabilities for all avian Level 3 classes given the true Level 3 class is black scoter. This completes the model, which can be visualized as a directed acyclic graph in Figure A1 in Appendix A. All model terms are listed and defined in (Table 2).

**Table 2.** Definitions of model terms by submodel and type. Submodels include: ecological, censoring (censor), and classification (class). Classification is further split into terms related to the survey data, validation data (valid), or both (shared). Term types are parameter (P), latent variable (LV), and data (D).

| Submodel | Type | Term | Definition |
| --- | --- | --- | --- |
| Ecological | P | $\beta_i = \{\beta_{0,i}, \beta_{1,i}\}$ | Expected abundance parameters of Level 3 class $i$ |
| Ecological | P | $\phi_i$ | Dispersion parameter of Level 3 class $i$ |
| Ecological | D | $x_j$ | Survey covariate value of image $j$ |
| Ecological | P | $\lambda_{i,j}$ | Expected abundance of Level 3 class $i$ in image $j$ |
| Ecological | P | $p_{i,j}$ | Abundance (NegBin) size parameter for Level 3 class $i$ in image $j$ |
| Ecological | LV | $y_{i,j}^{L3}$ | Abundance of Level 3 class $i$ in image $j$ |
| Ecological | LV | $y_{a,j}^{L1}$ | Abundance of Level 1 class $a$ in image $j$ |
| Censor | P | $\xi_a = \{\xi_{0,a}, \xi_{1,a}\}$ | Censoring process parameters of Level 1 class $a$ |
| Censor | P | $p_{a,j}^{retain}$ | Probability the censoring threshold is exceeded by class $a$ in image $j$ |
| Censor | D | $z_{a,j}$ | Binary censoring outcome of Level 1 class $a$ in image $j$ |
| Class (survey) | LV | $\gamma_l^s$ | Level 3 class of survey object $l$ |
| Class (survey) | D | $c_l^s$ | CNN Level 3 classification of survey object $l$ |
| Class (shared) | P | $\theta_{i,i'}^C$ | Probability a Level 3 object of class $i'$ is classified as class $i$ by CNN |
| Class (valid) | P | $\pi_i^v$ | Frequency of Level 3 class $i$ in validation data |
| Class (valid) | P | $\theta_{i,i'}^H$ | Probability a Level 3 object of class $i'$ is classified as class $i$ by Human |
| Class (valid) | LV | $\gamma_m^v$ | Level 3 class of validation object $m$ |
| Class (valid) | D | $c_m^v$ | CNN Level 3 classification of validation object $m$ |
| Class (valid) | D | $h_{m,1:3,o}^v$ | Hierarchical classification of validation object $m$ by human observer $o$ |

### Methods – Parameter Estimation

We jointly estimated the model parameters and latent variables using Markov chain Monte Carlo (MCMC) in the NIMBLE (de Valpine et al., 2017) package Version 0.12.1 for program R (R Core Team, 2021). We used uninformative priors except for ***θ***^*c*^ and ***θ***^*h*^ for which we used moderately informative priors to avoid posterior multi-modality. See Appendix A for a full description of prior distributions and details of two required custom MCMC updates.

### Methods – Simulation Study 1: Model Evaluation

We conducted two simulation studies of our model. First, we conducted a simulation study to assess how well the model parameters and the realized (but imperfectly observed) counts of Level 3 objects in images, ***Y***^L3^, can be estimated under a hypothetical scenario of ecological and sampling conditions. Further, we were interested in comparing the precision of parameter estimates and respective run times of the model with and without the image censoring process. We refer to these scenarios as S1a (with censoring) and S2a (without censoring), respectively.

We considered that there were 1000 survey images, 10 Level 3 object classes with the hierarchical labels delineated in Table 1, and that the expected abundance of all Level 3 classes except human-made responded to a single continuous covariate generated from a standard Normal distribution for each data set, whereas the expected abundance of the human-made class was assumed to be constant across images. The ecological parameters for abundance were chosen such that all avian, non-avian wildlife, and human-made objects had the same parameter values, respectively. We set *β*_0,*i*_ = −1 and *β*_1,*i*_ = 1.5 for avian species (*i* = 1,…, 5), *β*_0,*i*_ = −1.25 and *β*_1,*i*_ = 1 for non-avian species (*i* = 6,…, 9), and *β*_0,*i*_ = 0.5 and *β*_1,*i*_ = 0 for human-made objects (*i* = 10). We set the count dispersion parameter *ϕ* to 0.5 for all classes, indicating relatively large overdispersion compared to the Poisson distribution.

For the image censoring process (only included in S1a), we considered that the censoring decision was made only with respect to the number of avian objects in an image, setting *ξ*_0,*a*_ = 0 and *ξ*_1,*a*_ = −2.6 for *a* = 1, the avian Level 1 class. We set *ξ*_0,*a*_ = −8 and *ξ*_1,*a*_ = 0 for *a* ∈ (2, 3), the non-avian wildlife and human-made classes, yielding a 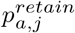 of effectively 0 with no response to the number of objects in an image. To improve estimation of the censoring parameters, we retained 15% of images with a censored outcome (*z_a,j_* = 0 for *a* = 1) at random, which were subjected to virtual CNN classification along with the uncensored images. These settings led to an average of 60% of images being discarded, with an average of 1.15 avian objects in censored images and 12.87 avian objects in retained images. For Scenario S2a, the image censoring process was not included.

We considered that both the human and CNN correct classification probabilities were 0.9 for all 10 Level 3 classes and that all incorrect classifications were equally likely. We considered there were 2 human observers who classified 200 validation objects of each Level 3 class and that these objects were also classified by the CNN. Simulated humans were able to classify objects to Level 1 with probability 0.99. Then, if a human made a Level 1 classification, they could classify objects to Level 2 with probability 0.95. Similarly, if a human made a Level 2 classification, they could make a Level 3 classification with probability 0.8. Unconditionally, humans could classify to Level 2 with probability 0.94 and to Level 3 with probability 0.75.

We fit our each model (with and without censoring) to 120 simulated data sets, running 3 chains with 15,000 iterations per chain. We thinned the structural parameters by 4 and the latent variables ***Y***^*L*3^ by 40. We used the posterior mode for point estimates and 95% highest posterior density (HPD) intervals for interval estimates. We assessed relative bias and coverage for all parameters, though for brevity, we present the mean bias and coverage for each parameter type averaged across Level 1 classes.

### Methods – Simulation Study 2: The Importance of Hierarchical Modeling for Modeling Misclassification

We conducted a second simulation study to demonstrate the importance of using a hierarchical model to estimate both ***Y***^*L*3^ and the ecological model parameters jointly. One of the key features of our model is that the ecological model determines the relative frequency that each Level 3 class occurs across images and this is vital information for estimating ***Y***^*L*3^. In fact, if we assumed 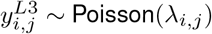, then this induces a prior on the Level 3 class of object *l* occurring in image *j* that is 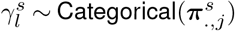, where 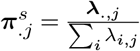 (Kéry and Royle, 2020). This same information is contained in other count models like the Negative Binomial that we use, but it cannot be summarized at the sample-level like it can for the Poisson (See Appendix B). Regardless, any count model used for the ecological process here carries the information about the relative frequencies of Level 3 objects across images.

In this second simulation study, we fit two simplified models to the same data sets that we simulated above for Scenario S2a (no image censoring). For the first simplified model, Scenario S2b, we considered the case where we only model the Level 3-level frequencies in a hierarchical model, ignoring how they vary across images. To do this, we assumed 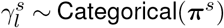, where ***π***^*s*^ is the frequency of Level 3 objects in the survey data set (note ***π***^*s*^ no longer varies across images). This is analogous to how we consider the relative frequencies in the validation data set with no image structure. In this model, 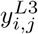 is a derived variable from each Level 3 object class, 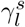, and image number, *ρ_l_*. We use this scenario to demonstrate that this model can estimate the overall number of Level 3 objects well, but estimates at the image-level will be biased because this model assumes the relative frequencies of Level 3 objects do not vary across images.

The most simple model we consider in Scenario S2c, is one where we simply ignore the relative frequencies of Level 3 objects and treat the Level 3 classes for each object as missing data to be updated from the model 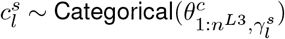. As for Scenario S2b, 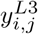 is computed as a derived variable. We use this scenario to demonstrate that this model cannot estimate the overall or image-level counts of Level 3 objects because it implicitly assumes each class occurs equally as often in the survey data and we simulated them to differ as would occur in real world situations.

## Results

### Results – Simulation Study 1: Model Evaluation

We found the parameter estimates (averaged over species) in Scenarios S1a and S2a to be roughly unbiased (< 1%) except for the negative binomial dispersion parameter, *ϕ*, which was positively biased by 5.79% with image censoring and 4.01% without (Table 3). This bias was more pronounced for rare species with the mean bias for avian objects being 3.4%, the mean bias for non-avian wildlife objects being 9.7% and the mean bias for human-made objects being 1.8% in Scenario S1a. The corresponding numbers for Scenario S2b are 3.6%, 5.5%, and 0.3%. For comparison, each of the 5 avian Level 3 classes had expected frequencies in simulated data sets of 0.123, each of the 4 non-avian wildlife classes had expected frequencies of 0.051 and the one human-made class had an expected frequency of 0.181 (Table 3, row 1).

**Table 3.** Percent relative bias of posterior modes, 95% coverage of highest posterior density (HPD) intervals, and mean posterior standard deviations for Scenarios S1a (with image censoring) and S2a (without image censoring). Results are averaged across all 10 Level 3 classes for each parameter type. *π^v^* is the species frequencies in the validation sample, *β*_0_ and *β*_1_ are the ecological process model regression parameters, *ξ*_0_ and *ξ*_1_ are the censoring parameters, Θ^*C*^ is the CNN classification probability matrix and Θ^*H*^ is the human classification probability matrix. We only assess the correct classification probabilities along the diagonal, i.e. 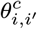 for all *i* = *i*′.

| Parm | % Relative Bias |  | Coverage |  | Mean SD |  |
| --- | --- | --- | --- | --- | --- | --- |
|  | S1a | S2a | S1a | S2a | S1a | S2a |
| $\beta_0$ | -0.60 | -0.43 | 0.946 | 0.941 | 0.148 | 0.100 |
| $\beta_1$ | 0.09 | -0.19 | 0.947 | 0.944 | 0.120 | 0.090 |
| $\phi$ | 5.79 | 4.01 | 0.957 | 0.947 | 0.109 | 0.086 |
| $\xi_0$ | -0.29 | . | 0.942 | . | 0.169 | . |
| $\xi_1$ | 0.15 | . | 0.917 | . | 0.064 | . |
| $\theta_{i,i'}^C$ | 0.81 | 0.64 | 0.917 | 0.893 | 0.023 | 0.022 |
| $\theta_{i,i'}^H$ | -0.45 | -0.43 | 0.925 | 0.921 | 0.018 | 0.018 |
| $\pi^v$ | -0.40 | -0.42 | 1.00 | 1.00 | 0.007 | 0.007 |

The coverage of the interval estimates were near nominal for each parameter type in both models S1a and S2a, though there may have been reduced coverage for the classification probability matrix parameters due to the moderately informative prior. Scenario S2a without image censoring estimated the ecological model parameters more precisely as judged by the mean posterior standard deviations (Table 3) than the model with censoring, and the CNN classification probabilities were estimated slightly more precisely. The mean per chain run time were 81.35 and 88.70 minutes for Scenarios S1a and S2a, respectively, corresponding to the censored version being 8% faster.

### Results – Simulation Study 2: The Importance of Hierarchical Modeling for Modeling Misclassification

In the second simulation study, we found that the models in Scenarios S2a and S2b estimate the overall number of Level 3 objects with less than 1% bias except for the human-made class in Scenario S2a, with a relative bias of −1.97%. The 95% coverage was near nominal for these two Scenarios and Scenario S2a that included the ecological process model estimated the overall number of Level 3 objects more precisely as judged by the coefficient of variation (CV; Table 4). The model in Scenario S2c underestimated the total number of objects in the most common classes (Avian and human-made) and overestimated the total number of non-avian wildlife objects (Table 4).

**Table 4.** Comparison of bias, 95% coverage, and coefficient of variation (CV) of Total L3 and L3 x image counts across Scenarios S2a (ecologically-adjusted), S2b (frequency adjusted), and S2c (unadjusted). These comparisons are made across the L1 classes, avian, non-avian wildlife (NAW) and human-made, with each metric averaged across the L3 classes in each L1 class. Frequency is the expected proportion of the counts for each L3 group. For total L3 counts, we present relative bias and for L3 x image counts, we present absolute bias (due to many true 0’s).

| Count Type | Metric | Scen | L1 Class |  |  |
| --- | --- | --- | --- | --- | --- |
|  |  |  | Avian | NAW | human-made |
| Total L3 | Frequency | . | 0.123 | 0.051 | 0.181 |
| Total L3 | RelBias | *S2a* | -0.16 | 0.70 | -1.97 |
| Total L3 | RelBias | *S2b* | 0.270 | -0.39 | 0.21 |
| Total L3 | RelBias | *S2c* | -4.86 | 10.45 | -17.52 |
| Total L3 | Cover | *S2a* | 0.98 | 0.98 | 0.97 |
| Total L3 | Cover | *S2b* | 0.96 | 0.97 | 0.91 |
| Total L3 | Cover | *S2c* | 0.96 | 0.98 | 0.89 |
| Total L3 | CV | *S2a* | 3.35 | 5.91 | 2.05 |
| Total L3 | CV | *S2b* | 4.24 | 7.83 | 3.35 |
| Total L3 | CV | *S2c* | 3.27 | 5.106 | 2.71 |
| L3 x image | AbsBias | *S2a* | -0.11 | 1.41 | -1.07 |
| L3 x image | AbsBias | *S2b* | 0.21 | -0.70 | 0.13 |
| L3 x image | AbsBias | *S2c* | -3.62 | 20.24 | -9.51 |
| L3 x image | Cover | *S2a* | 0.93 | 0.93 | 0.87 |
| L3 x image | Cover | *S2b* | 0.94 | 0.93 | 0.89 |
| L3 x image | Cover | *S2c* | 0.73 | 0.08 | 0.05 |

For the estimates of Level 3 class by image counts the absolute bias in Scenarios S2a and S2b was minimal (Table 4), though there was more bias in Scenario S2a that overestimated the number of each non-avian wildlife class by 1.41 counts and underestimated human-made by 1.07 counts (Table 4). Coverage was nominal for Scenarios S2a and S2b, except for the human-made class, which was marginally too low (Table 4). Scenario S2c produced biased Level 3 class by image count estimates with very low coverage, particularly for non-avian wildlife and human-made objects (Table 4).

While the models in Scenario S2a and S2b appeared to both perform well when assessing the bias and coverage of the total Level 3 counts and Level 3 class by image counts, the deficiency of the model in Scenario S2b appears when plotting the estimates as a function of the number of objects of each type in the images (Figure 2). We see that Model S2b estimates the total number of Level 3 counts well, but for the Level 3 by image counts the model underestimates the largest counts and overestimates the smallest counts. Thus, the negative bias for larger counts and the positive bias for smaller counts roughly cancels out and the poor coverage for higher counts is offset by greater than nominal coverage for smaller counts. Only the model in Scenario S2a estimates the counts of all sizes well.

**Figure 2.**
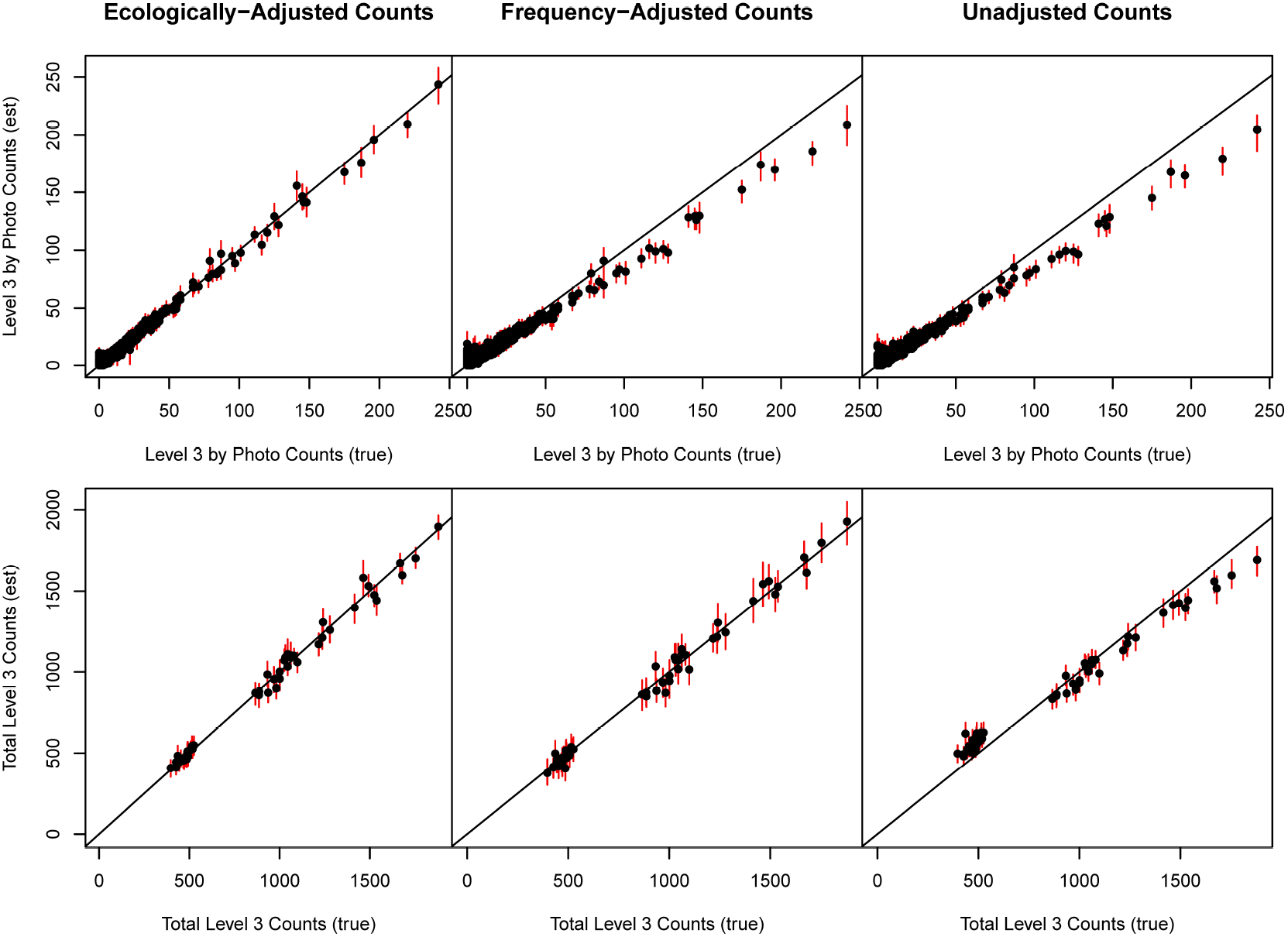
Simulation Study 2 Results. Estimated vs. true overall Level 3 counts and Level 3 by image counts for Scenarios S2a (ecologically-adjusted), S2b (frequency-adjusted), and S2c (unadjusted). Black dots are posterior means and red lines are 95% highsets posterior density (HPD) intervals.

## Discussion

We developed a hierarchical model that allows for the estimation of species-specific abundance from aerial images where object detection and (imperfect) classification are done by deep learning and images unlikely to contain targeted species are censored, leading to preferential sampling. This model is tailored to the ongoing USFWS/BOEM aerial survey and deep learning data collection and processing methods for marine wildlife. Specifically, the model accommodates both the deep learning classification at multiple taxonomic levels and hierarchical human labeling process to produce validation data, which is subject to missing data and validation errors at all taxonomic levels. Further, we demonstrated the utility of coupling the ecological model describing how the objects to be classified are observed with the model(s) for taxonomic classification for probabilistically resolving the true classes and making ecological inference.

Our model performed well at the simulation settings we considered, which included a particularly challenging level of overdispersion in Level 3 counts and a correct classification probability of 0.90 by both the humans and CNN for all species. We did find that the estimates of the overdispersion parameters, ***ϕ***, were positively biased; however all other parameters were well estimated in the presence of this bias, and the coverage for ***ϕ*** was roughly nominal. The bias in ***ϕ*** was largest for Level 3 classes with the lowest total abundance in the data sets and we expect that the bias can be largely removed by collecting more than the 1000 images we considered, at least when the CNN correct classification probability is as high as 0.9.

Under our simulation settings, the drawbacks of image censoring may outweigh the benefits from the perspective of the computation time to fit the statistical model. Discarding images reduced the precision of the ecological model parameter estimates, and the Level 3 by image counts were necessarily estimated less precisely when reducing the information from object-level classifications to image-level censoring outcomes. The benefit of image censoring in our simulations was a modest decrease in computation time (≈ 8%). Therefore, the decision regarding image censoring may be most influenced by the additional time required to subject these images to Level 3 classification by the CNN, though the decrease in computation time will be greater in systems with more objects that are not of interest than we considered in Simulation Study 1. Further, more information could be retained from images not subjected to Level 3 CNN classification than we considered. If instead, we retained the object-level Level 1 classifications, we could model them in the same manner that we model Level 3 classifications–there is some information about the Level 3 class of all objects in an image in the image level censoring outcome, but there is more information about the Level 3 class of each object in the Level 1 classification for that object.

While this model has many of the important features of aerial surveys for marine wildlife conducted by USFWS/BOEM, we make some assumptions that may be difficult to meet in practice. First, we assume the CNN is perfectly transferable between the validation data set for which humans provide classifications, and the survey data to be classified without the aid of humans. By “perfectly transferable”, we mean ***θ***_*C*_ is exactly the same for the validation and survey data sets. If transferability is a concern; however, the validation images can be selected randomly from the survey images themselves (see Spiers et al., 2022, for an example), though this requires new human validation data to be produced for every survey. Further, we did not consider that there may be image-level variability in ***θ***^*C*^, perhaps as a function of image quality or camera distance from the water surface. We expect unmodeled heterogeneity in ***θ***^*C*^ to erode the reliability of ecological inference, but this heterogeneity can be accommodated with fixed or random image effects (which also must be “perfectly transferable” if the validation images are from a different survey). This approach would be similar to the use of sample quality covariates for genotyping error rates in capture-recapture models (Augustine et al., 2020)–as not all genetic samples have the same genotyping error rates, not all images yield the same Level 3 class error rates.

A second set of assumptions that may be challenging to meet are assumptions of classifier independence. For the validation data, we assume the human observer classifications are independent of one another and independent of the CNN classifications. Then for the survey data, we assume the CNN classifications are independent of the censoring outcomes (or higher-order classifications, if retained). Each of these sets of random variables may be correlated due to several factors, with perhaps the most likely source being features relating to the quality of the images. With respect to the validation data, if the main factor determining whether humans and the CNN can correctly classify the species is image quality, then humans and the CNN will tend to make more of the same correct and incorrect scores than expected under independence. This correlation in classification errors could potentially be modeled, with image-level covariates or correlated random effects, but 1) more human validation is required as the correlation increases and 2) if this correlation is near 1.0, we effectively have only one observer and this type of model will not work as intended.

A third challenging assumption is that the human hierarchical classifications are missing at random. Presumably, some objects are easier to classify to Level 3 than others and this may be due to features of the images or of features of the Level 3 class. If classifications are missing as a function of image quality, the set of human classifications that are not missing are the ones from the best images where both the humans and CNN are more likely to make correct classifications. Then, these correct classification probabilities estimated from the best images will be applied to all images, overestimating the correct classification probabilities for the poor images and eroding the reliability of ecological inference. If this effect is only present for particular species, one possible solution is to define their Level 3 class to be a higher order, such as genus, if genus can be more reliably identified than species. Another possible remedy may be to change the survey design, for example, perhaps the plane could fly lower or better cameras could be used (Hong et al., 2019). Ultimately, if humans cannot identify a substantial subset of objects in images, we cannot accurately quantify the performance of the CNN and correct for misidentification error. In this case, we require that the CNN is near perfect for all Level 3 classes for reliable ecological inference; however this is difficult to demonstrate without knowing the true Level 3 classes of the validation samples.

A final factor of this survey design that we did not consider is that the object detection will not be perfect (Hong et al., 2019). Deep learning object detection algorithms such as YOLO generate boxes around objects of interest and provide a classification for the object inside the box. It is possible that objects can be missed entirely, boxes may be drawn around more than one object, or multiple boxes are drawn for the same object. Our hierarchical model could potentially be expanded to accommodate these types of errors with additional types of human validation data, or these error rates may be modeled separately and applied to ***Y***^*L*3^ to produce the final estimates as derived variables. In either case, we feel this is not a trivial problem particularly for single objects assigned two boxes or multiple objects, potentially of different Level 3 classes, assigned a single box. These “erroneous objects” will be assigned single Level 3 classes by the CNN, but the estimated CNN classification probabilities from the human validation data in the current survey protocol are for correctly identified objects only (1 box per object). Therefore, the current model not only does not correct for detection errors; its ability to correct for object misclassification will also be eroded by these “erroneous objects” to the extent that they occur in practice.

Our second simulation study demonstrates the importance of considering the relative frequencies of each Level 3 class (or species more generally) to the task of classification. From the standpoint of the classification model, the ecological model parameters induce a prior distribution on class membership to be considered along with the classification model (Kéry and Royle, 2020). As mentioned above, when the count model is Poisson, this prior can be described at the sample level. For other count models, this prior information cannot be disaggregated to the sample level, but the prior information about relative frequency is still contained in the count model. We demonstrated that the information about the relative frequency of classes needs to be considered in conjunction with the classification model at the level the ecological model will be modeled for unbiased inference. In our simulations, modeling the Level 3 class frequencies was sufficient for estimating the Level 3 counts across images, but the Level 3 by image count estimates were biased for images with very few or very many objects (Figure 2). When the ecological data are structured by image (or more generally space), we require a model for the relative frequencies of Level 3 objects at the image-level. Similarly, for an ecological model with replication across occasions, such as an occupancy model, we require a model for the relative Level 3 class frequencies across space and time. The most efficient model for this task is the ecological model that produced the observations (or our best approximation of it) and combining the ecological and classification models into a single hierarchical model allows all parameters to be estimated jointly with uncertainty propagated appropriately (Kéry and Royle, 2020). However, this approach does come at a cost–because the estimated true classes depend on the ecological model structure, more care is required to specify it correctly than when all Level 3 class identities are known with certainty. We expect that the sensitivity to ecological model misspecification increases as the correct classification probabilities decrease.

In summary, we have described a general framework for estimating marine wildlife abundance from aerial images for the survey methodology in use by USFWS/BOEM. In doing so, we outlined the assumptions that are required for this approach and how they may not be met. Thus, our model identifies areas for further investigation and provides a means to assess the effects of violating these difficult assumptions once data are in hand. Further, our model can provide a starting framework for an expanded model accounting for detection errors. Secondly, we demonstrated ways in which the ecological model that produced the observations to be classified is relevant to the classification task itself, and this has implications for how we make ecological inference using CNN-classified observations. Often, the classification and ecological modeling tasks are considered separately, but we argue they are fundamentally linked and recognizing this fact can improve ecological inference and provide a means to assess what classification accuracy is required (for every class) for particular ecological systems and models via simulation studies. To date, most of the focus in deep learning applications to wildlife photography has been on improving algorithms to increase their accuracy and precision, but the performance achieved is difficult to interpret without knowing the performance required for reliable ecological inference.

## Acknowledgements

We thank the Bureau of Ocean Energy Management and, in particular, Timothy White, for collaboration in integration of remote sensing and machine learning technologies for marine wildlife surveys. We also acknowledge staff of the U.S. Geological Survey’s Upper Midwest Ecological Sciences Center for developing annotation tools and managing the process training and validation data development for this project. Finally, we thank Paul Conn and Josh Twining for constructive feedback on a previous version of this manuscript. Some technical developments that contributed to this work resulted from the Working Group on Coupled Classification funded by the USGS John Wesley Powell Center for Analysis and Synthesis. The findings and conclusions in this article are those of the author(s) and do not necessarily represent the views of the U.S. Fish and Wildlife Service. Any use of trade, firm, or product names is for descriptive purposes only and does not imply endorsement by the U.S. Government.

## SI Appendices

## Appendix A Directed Acyclic Graph (DAG), Prior Distributions, and MCMC Details

## Model DAG

**Figure A1.**
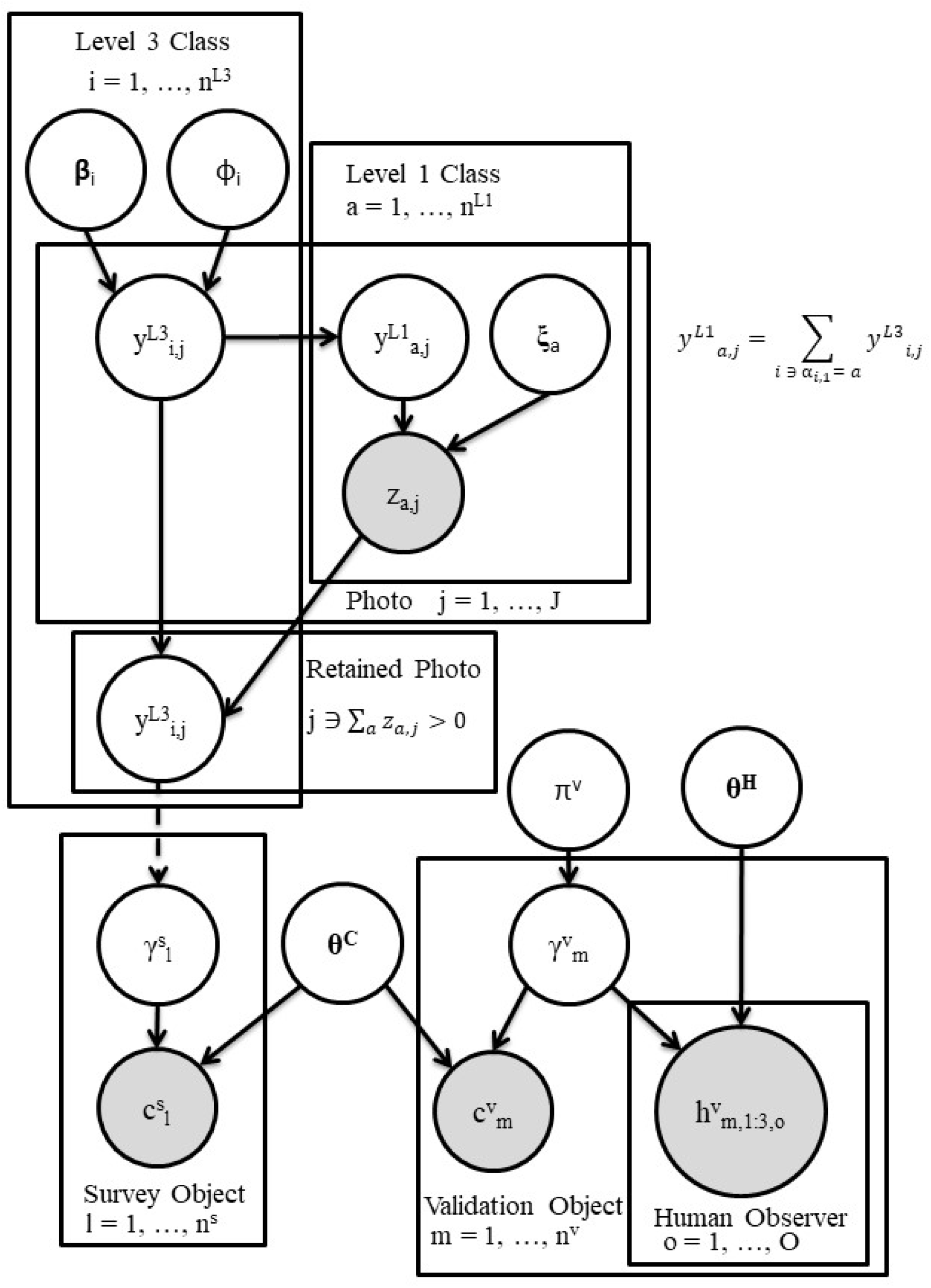
Directed Acyclic Graph of the hierarchical model. (Provide some level of description here. Observed nodes have a grey fill, though the human classification may only be partially observed.

## Prior Distributions

We use uninformative to weakly informative priors for all model parameters except for the CNN and human classification probability matrix where we place a roughly uniform prior on the probability of correct classification, which removed multimodality from the posterior distributions for some data sets (Figure A1).

- Validation data species proportions

– 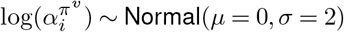
– ***π***^*v*^ ~ Dirichlet(***α***^π^)
- CNN classification probability matrix for true Level 3 class *i*

– 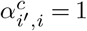 for *i*′ = *i*, 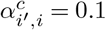 for *i*′ ≠ *i*
– 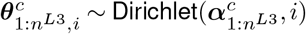
- Human classification probability matrix for true Level 3 class *i*

– 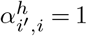 for *i*′ = *i*, 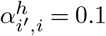 for *i*′ ≠ *i*
– 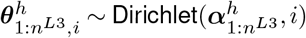
- Ecological process model parameters

– *ϕ_i_* ~ Uniform(0, 25)
– *β*_0,*i*_ ~ Normal(*μ* = 0, *σ* = 10)
– *β*_1,*i*_ ~ Normal(*μ* = 0, *σ* = 10)
- image censoring process parameters

– *ξ*_0,*i*_ ~ Logistic(*μ* = 0, *τ* = 1) (*τ* is the rate)
– *ξ*_1,*i*_ ~ Normal(*μ* = 0, *σ* = 10)

## MCMC Details

We use default MCMC samplers supplied by Nimble except for updating 1) the latent variables in discarded images and 2) the latent variables in retained images. We use the following algorithm:

(**0**) Initialize the latent variables. First, we initialize the latent variables for retained survey images. We set the true Level 3 class, 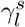 to be the CNN Level 3 predicted class, 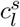 for objects whose image number *ρ_l_* is in the set of retained images. Then, the initialized true Level 3 classes determine the initial values for the *j* indices of 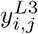. Second, we treat the initialized Level 3 by image counts for retained survey images as truth and estimate the ecological model parameters, ***β***_0_, ***β***_1_, and ***ϕ***, using negative binomial regression independently for each Level 3 class. We set the initial values of the ecological model parameters, 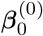, 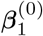, and ***ϕ***^(0)^ to these estimates from a possibly true configuration of the Level 3 by image counts for retained images. Next, we simulate the Level 3 by image counts for discarded survey images from the initial ecological parameters, 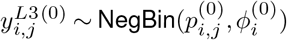. Then, we calculate the initial Level 1 by image counts, ***y***^*L*1(0)^, implied by the initial Level 3 counts, ***y***^*L*3(0)^. Finally, we use the first human observer to initialize the true Level 3 class of object *m*, setting 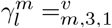 when 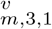 is observed. When 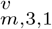 is not observed, we draw 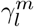 at random from the classes consistent with the lowest level of taxonomic accuracy provided by the human.

(**1**) Update the latent variables in discarded images, ***y***^*L*3^, and ***y***^*L*1^. For every discarded image, *j*, we update the latent variables one Level 3 class at a time using a Metropolis-Hastings step. The full conditional of 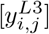 is proportional to the product of the likelihood of the ecological process model and the censoring model

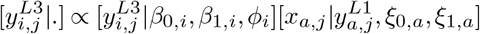

where

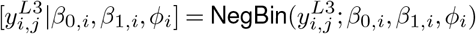

and

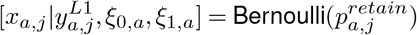

**Figure A2.**
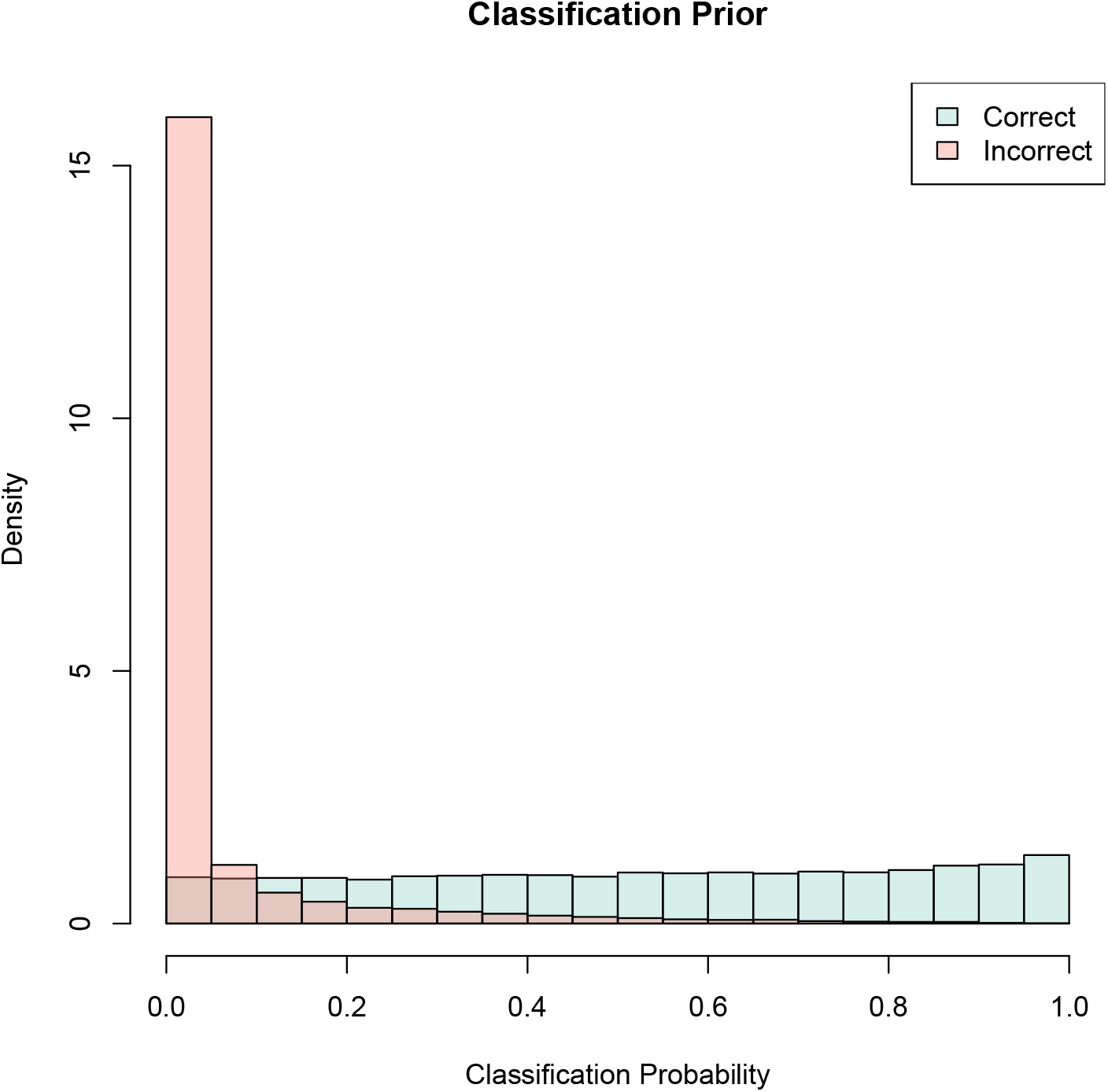
Simulated data from the prior distributions for the CNN and human classification probability matrices. Correct and incorrect classification events are when *i*′ = *i* (along the diagonal) and *i*′ ≠ *i* (off-diagonal), respectively.

First, we generate a proposal 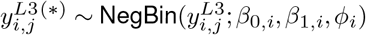 which implies the proposal 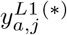. Then, we compute 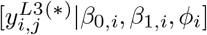 and 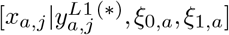. The Metropolis-Hastings acceptance probability is then

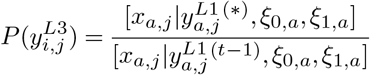

where *t* is the iteration number. Note, because we use 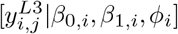 as a proposal and this is the same as the likelihood, they cancel out.

(**2**) Update the latent variables in retained images, ***γ***^*s*^, ***y***^*L*3^, and ***y***^*L*3^. For every retained image *j*, we update each 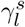 whose *ρ_l_* = *j* one at a time. The full conditional of 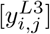 is proportional to the product of the likelihood of the ecological process model, the classification model, and the censoring model

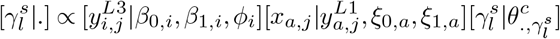

where

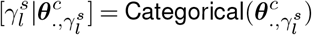

For each survey object *l* located in image *ρ_l_* = *j*, we generate a proposal 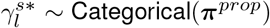 where 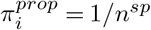 for *i* = 1,…, *n^sp^*, which is symmetric. Then, we compute 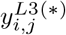 and 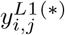 using 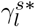, and compute 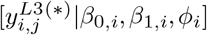 and 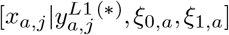. The Metropolis-Hastings acceptance probability is then

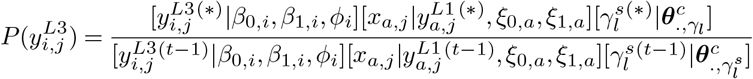

## Appendix B Simulation Study 3

We conducted a third simulation study to demonstrate how the information about the Level 3 class contained in the ecological process model varies as a function of the count model dispersion, *ϕ*. First, note that Simulation Study 2 demonstrated the utility of using the ecological model as a prior for the Level 3 class frequencies in the survey data. These frequencies are not structural parameters of our model, though they could be used in that manner if the counts are Poisson (Kéry and Royle, 2020; Spiers et al., 2022). As noted in the main text, if 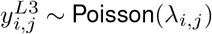, then for object *l* in image *j*, 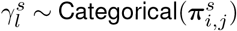, where 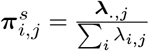 (Kéry and Royle, 2020). Thus, 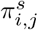 is the expected probability a randomly selected object in image *j* will be of Level 3 class *i*.

We propose that one measure of the information content about the Level 3 class of each object contained in the ecological model is the variance of the realized Level 3 class frequencies, 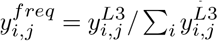 for each Level 3 class across images when the ecological count model parameters are known. A better measure would consider the uncertainty in the ecological model parameter estimates, but we can make some general points without that consideration. We used Monte Carlo simulation to generate the realized Level 3 class frequencies across images in a simplified set of scenarios where all ten species have the same parameters and do not respond to habitat covariates. We simulated Level 3 class by image data following 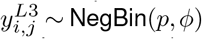, where 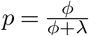. We considered six scenarios with all possible combinations of *λ* = 5 and 1.5 and *ϕ* = 5000 (approximately Poisson), 5, and 0.5 across 1 million images each. After simulating 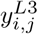, we calculated 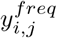 following the equation above. Finally, we plot the distributions of the realized Level 3 class frequencies to show that the variance grows with the level of overdispersion in the count model.

Figure B1 demonstrates that as we increase the variance in the ecological count model, we increase the variation in the realized Level 3 class frequencies across images. As a result, the ecological count model will be less informative of the Level 3 classes of objects. A further observation is that the distributions for the scenarios with *λ* = 0.15 and *ϕ* = 5 and 0.5 are bimodal with one mode at zero and a second mode between 0 and 1. We have found that using MCMC to fit “coupled classification” models (Kéry and Royle, 2020) with latent species or individual identities with the complete data likelihood as we use here can lead to non-convergence, biased estimation, or an underestimation of posterior variances due to complicated posterior geometry (Betancourt and Girolami, 2015). The bimodal distributions of 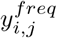 illustrate the source of this issue–the number of objects of Level 3 class *i* in image *j* may be 0 or some higher number, but the probability may be effectively 0 in between. If so, and you update the Level 3 class of one object at a time, you may not be able to traverse this low probability region and get stuck in one mode. This could potentially happen for many Level 3 classes in many images.

The effect of the classification model likelihood on the posterior of 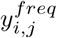 is not conveyed in Figure B1, but if the correct classification probability is high enough, this information may rule out one of the two modes in the distribution of 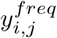. If this is the case for all images, estimation will be correct if 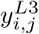 is initialized in these corresponding configurations, which may happen frequently when the correct classification probabilities are high (though not guaranteed) or 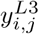 may be able to converge to the correct posteriors modes while the ecological count and classification model parameters are converging.

We did not observe any problems of convergence or sampling in our simulation studies; however, this was common when using a zero-inflated Poisson distribution instead of the negative binomial we focus on here. We attribute this to the fact that zero inflation introduces more 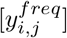 modes at 0 and the lack of overdispersion increases the chance these two regions of the posterior are traversable using MCMC updating the Level 3 class of one object at a time.

We see two possible solutions to this problem. First, Level 3 class can be marginalized, which was the approach taken by Wright et al. (2020) in a coupled classification occupancy model (where Level 3 class was “species” and the spatial units were “sites”). This approach is not workable for a count model other than the Poisson because the Poisson is the only count distribution where the individual count members have independent distributions and we do not need to sum over all possible configurations of Level 3 classes to objects in an image, which can be computationally intractable. Wright et al. (2020) allow for extra-Poisson variation via random effects and bypass this issue. A second challenge is that if there is zero inflation, the only way to specify a model that allows for marginalizing Level 3 classes is to condition Poisson counts on binary zero-inflation indicators. For example, Wright et al. (2020) specify *z_i,j_* to be occupancy states and specify [*y_i,j,k_| λ_i,j,k_, z_i,j_*] ~ Poisson(*λ_i,j,k_* × *z_i,j_*) for species *i* at site *j* on occasion *k* (using notation closer to that used in this paper). Then, for each site, they can marginalize over all possible combinations of ***z***_.,*j*_ and all possible combinations of sample-level Level 3 classes, 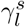 for samples collected at site *j*. This allows the Level 3 class to be fully marginalized, removing the MCMC challenges identified above. Unfortunately, this approach is species-limited because the number of all possible ***z***_.,*j*_ states is 2 raised to the number of species, which grows exponentially as a function of the number of species. For example Wright et al. (2020) considered 8 species with 256 possible combinations. Raising this to the 10 Level 3 classes in our study yields 1024 possible combinations. For Spiers et al. (2022) with 62 species, there are over 4.6 × 10^18^ possible combinations, which would be computationally intractable. Finally, Rhinehart et al. (2022), which we include in the category of “coupled classification” occupancy models, only considered 2 Level 3 classes (focal species vs. other), but future users should be wary about MCMC sampling challenges when adding more species.

A second solution to poor MCMC performance is to propose to update all Level 3 classes in an image in a joint Metropolis-Hastings update. If there are no CNN classification data associated with each image and counts are Poisson, 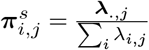 is a sensible proposal joint proposal distribution (in fact, it is the full conditional distribution; Chandler and Royle, 2013). However, when we introduce CNN classification data associated with each image, the proposals from this distribution will almost always be rejected, particularly when the CNN correct classification probabilities are high and when there are more objects in each image (personal observation). Thus, we opted to update the Level 3 class of one object at a time, where the CNN score of each object can be considered in the proposal. For this particular sea duck application a potential third solution would be to model the transects as the spatial unit instead of each image. The idea here is that fewer species may have a mode in 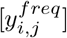 at zero when you sum their counts over all the images in a transect. However, the image censoring process occurs at the image level which would need to be modified, if possible, to decrease the modeled spatial resolution.

We recommend these issues be investigated further, and alternative MCMC strategies considered. In the mean time, we recommend marginalizing over species/Level 3 class when possible and thorough simulation investigations to assess convergence and mixing when it is not

**Figure B1.**
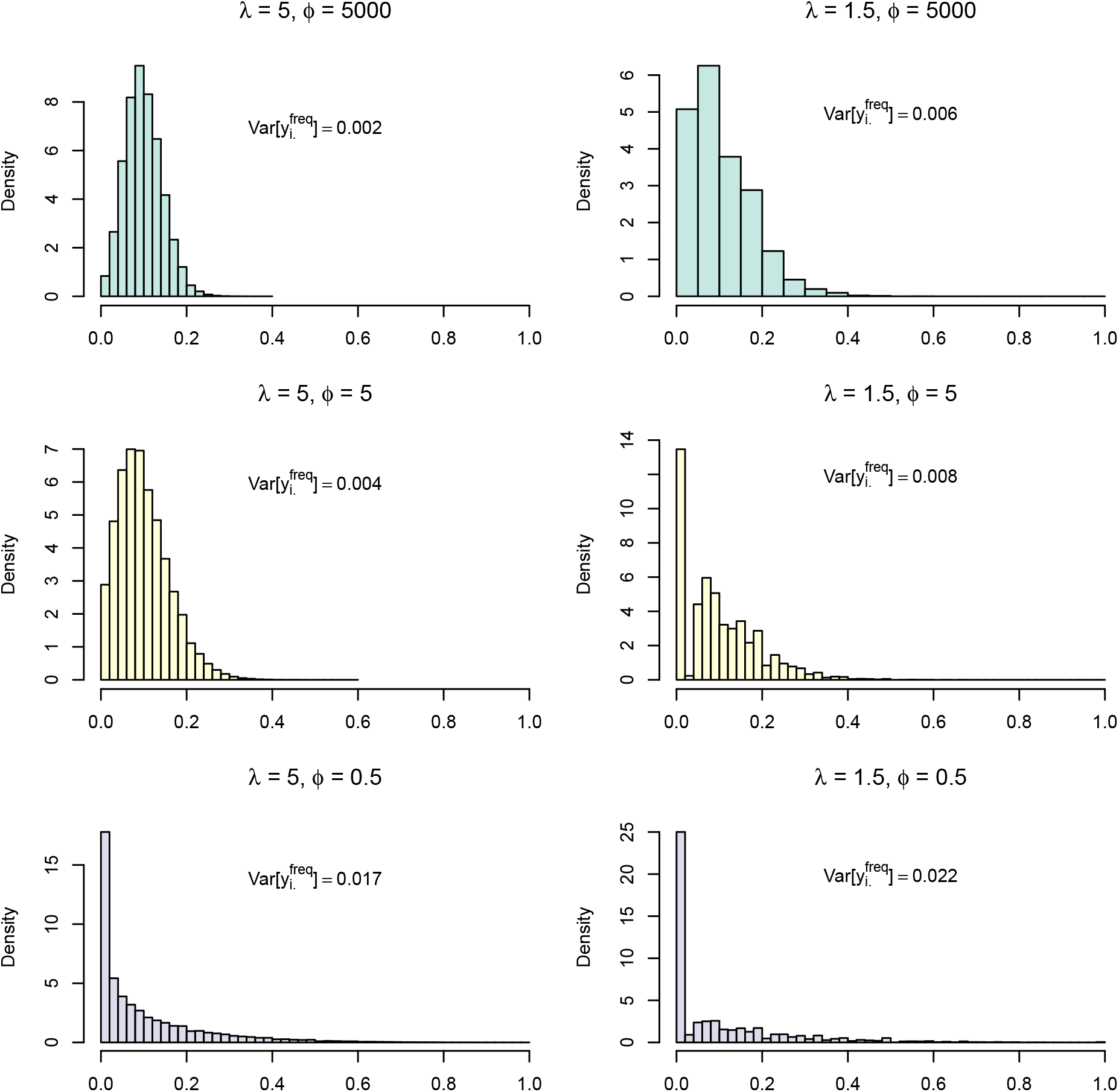
Distributions of realized species frequencies, 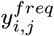, in each image across two levels of *λ* and three levels of *ϕ* from Simulation Study 3. Smaller values of *ϕ* indicate more dispersion and *ϕ* = 5000 leads to and approximately Poisson distribution. The variance of each scenario is printed on the plot and 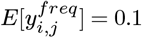 in all scenarios.

